# Small Molecule Inhibition of the Innate Immune Response Increases Transgene Expression

**DOI:** 10.1101/338707

**Authors:** Kyle Spivack, Christine Muzzelo, Christopher Neely, Julia Vanzelli, Evan Kurt, Jacob Elmer

## Abstract

Foreign molecules like plasmid DNA trigger a complex and potent innate immune response comprised of highly redundant signal transduction cascades that result in the activation of transcription factors and the production of inflammatory cytokines. Unfortunately, this defense mechanism can hinder gene therapy by inhibiting transgene expression. The goal of this study was to increase transgene expression by inhibiting key components of the innate immune response (β-catenin, NF-κB/AP1, TBK1, TLR9, and p38 MAPK) with small molecule inhibitors (iCRT-14, curcumin, BX-795, E6446, and VX-702 respectively). The effects of each drug on transgene (luciferase) expression, inflammatory cytokine (IL-6) levels, and cell viability were quantified in prostate (PC3), breast (MCF-7), and murine bladder (MB49) cancer cell lines. The β-catenin inhibitor iCRT-14 (1 μM) provided the highest enhancement of 35.5 ± 19-fold in MCF-7 cells, while the other inhibitors increased transgene expression at a more modest level (2-9 fold). The optimal concentrations of iCRT-14, curcumin, and VX-702 showed no significant effect on cell proliferation; however, optimal concentrations of BX-795 and E6446 did significantly reduce cell proliferation. Nonetheless, inhibition of the innate immune response by iCRT-14 and curcumin was confirmed by a concomitant decrease in IL-6 production in PC3 cells. These results demonstrate that these inhibitors can improve gene therapy by preventing an inflammatory innate immune response.

## Introduction

The innate immune response (IIR) is the cell’s first line of defense against infection.^1–3^ It is comprised of several enzymes and pathways which actively monitor the cell for signs of bacterial or viral infection. For example, cytosolic DNA is recognized as a sign of infection since host cell DNA is restricted to the nucleus.^4^ Once cytosolic DNA is detected by the IIR, it triggers a signaling cascade of kinases that ultimately results in the activation of transcription factors that induce the expression of inflammatory cytokines (e.g. IL-6 and TNF-α). Those cytokines can then activate additional genes that help defend the cell against the pathogen by inducing apoptosis, inhibiting protein translation, or other mechanisms.^5^ For example, when the cytosolic DNA sensor DAI (DNA-dependent activator of IRF) binds to DNA, it initiates a series of phosphorylation events involving IKK and TBK kinases, along with the transcription factors IRF3 and NF-κB. The activated transcription factors then translocate to the nucleus and upregulate expression of Type I interferons as shown in Figure 1.^6^ Another well studied pathway begins with Toll-Like Receptor 9 (TLR9), which specifically binds to dsDNA with unmethylated CpG motifs (CpGs in human DNA are methylated).^7–9^ TLR9 then recruits MyD88 to phosphorylate multiple kinases (IRAK 1, JNK, IKKα/β/γ), which in turn activate multiple transcription factors (AP-1, NF-κB, IRF-5, & IRF-7) to induce cytokine expression (e.g. IFNα).^10,11^

**Figure 1.**
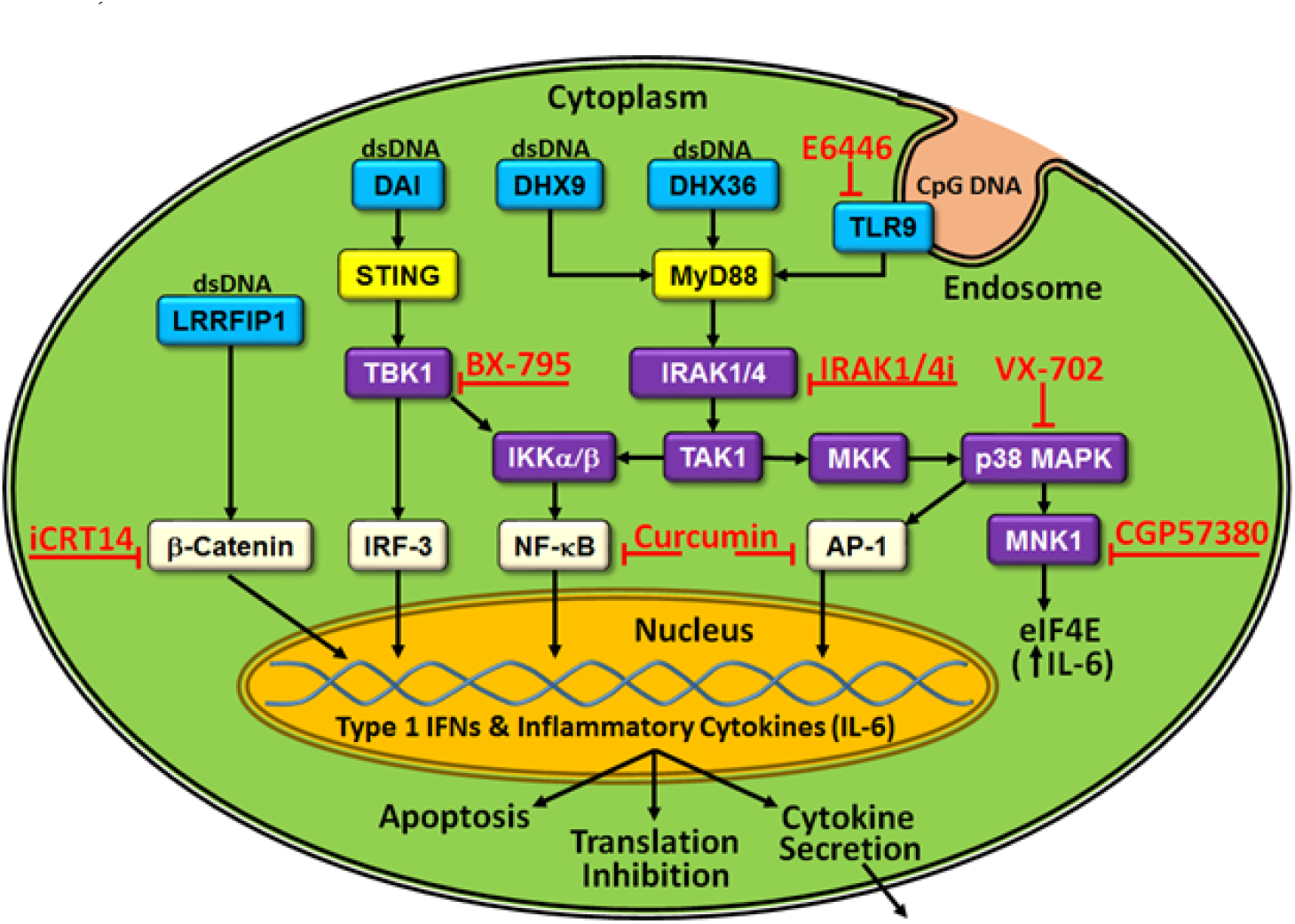
Examples of the innate immune response to cytosolic DNA. DNA sensors (blue) bind to DNA and subsequently (through yellow intermediates) activate kinases (purple). A series of phosphorylation events then activate transcription factors (white) that translocate to the nucleus to induce the expression of Type 1 IFNs and inflammatory cytokines which are secreted and propagate the innate immune response to other cells.

Unfortunately, while these pathways provide a crucial defense against foreign pathogens, they can also hinder gene therapy since most gene therapy techniques introduce plasmid DNA into the cytosol.^12–14^ Indeed, plasmid DNA has been shown to increase the expression of inflammatory cytokines, including IL-6 and TNF-α, which have been shown to decrease transgene expression.^4,15^ For example, Quin et al demonstrated that the addition of interferon-γ (IFN-γ) and TNF-α τo C2C12 myoblasts inhibits transgene expression.^16^ Consequently, the innate immune response must be carefully considered when developing techniques for gene therapy.

Many previous attempts to improve gene therapy via inhibition of the IIR have focused on TLR9.^17,18^ For example, since TLR9 is activated by unmethylated CpG motifs, Hyde et al removed all of the CpG motifs from a plasmid DNA sequence. The resulting “CpG-free” plasmid did not activate TLR9 and did not induce inflammation in mice.^18^ However, reintroducing a single CpG motif into the plasmid was sufficient to activate TLR9. Alternatively, *in-vitro* methylation of CpGs in plasmid DNA was also shown to prevent TLR9 activation and prolong transgene expression by up to four weeks when compared to unmethylated CpGs.^19^ In addition to TLR9, another recent study showed that inhibition of TBK-1 kinase with the small molecule inhibitor BX-795 enhanced lentiviral transgene expression in natural killer (NK92) cells by an average of 3.8-fold.^20^ Finally, silencing of the receptor for IFNα/γ (IFNAR) with shRNA has also been reported to enhance transgene expression.^21,22^

All of these previous studies clearly show that transgene expression can be improved by inhibiting different parts of the IIR. The goal of this work was to further improve gene therapy by targeting additional components of the IIR with the small molecule inhibitors (SMIs) shown in Figure 1.^23–27^ For example, the transcription factor β-catenin was inhibited with iCRT-14, while TLR9 was inhibited with the novel inhibitor E6446.^23,25,28^ The kinases TBK-1 and p38 MAPK were also inhibited with BX-795 and VX-702, respectively.^27^ Finally, curcumin was included as a general anti-inflammatory drug that inhibits cytokine production (TNF-α, IL-6, IL-12) by suppressing the activation of transcription factors NF-κB and AP-1 associated with the innate immune response.^24^ The effects of each drug on transgene (luciferase) and cytokine (IL-6) expression were tested.

## Materials and Methods

### Reagents and Materials

Polyplexes were prepared by mixing branched polyethyleneimine (PEI) from Sigma Aldrich (MW = 25,000) with a luciferase expression plasmid (pGL4.50) from Promega (Madison, WI). Small molecule inhibitors iCRT-14 {5-[[2,5-Dimethyl-1-(3-pyridinyl)-1H-pyrrol-3-yl]methylene]-3-phenyl-2,4-thiazolidinedione}, BX-795 {N-[3-[[5-iodo-4-[[3-[(2-thienylcarbonyl)amino]propyl]amino]-2-pyrimidinyl]amino]phenyl]-1-Pyrrolidinecarboxamide hydrochloride} and curcumin {(E,E)-1,7-bis(4-Hydroxy-3-methoxyphenyl)-1,6-heptadiene-3,5-dione} were purchased from Sigma Aldrich. VX-702 {6-[(aminocarbonyl)(2,6-difluorophenyl)amino]-2-(2,4-difluorophenyl)-3-pyridinecarboxamide} was purchased from Cayman Chemical (Ann Arbor, MI) and E6446{6-[3-(pyrrolidin-1-yl)propoxy)-2-(4-(3-(pyrrolidin-1-yl)propoxy)phenyl]benzo[d]oxazole} was generously provided by Eisai Inc (Tokyo, Japan). In addition, CGP 57380, an MNK1 inhibitor, was purchased from Cayman Chemical and IRAK1/4i (an IRAK 1 and 4 inhibitor) was purchased from Sigma Aldrich. These small molecule inhibitors were dissolved in DMSO and stored at −72°C when not in use.

PC3 cells were purchased from ATCC (Cat# CRL-1573) and MB49 cells were provided by Christina Voelkel-Johnson of the Medical University of South Carolina. PC3 and MB49 cells were cultured in RPMI-1640. MCF-7 cells were a generous gift from Dr. Janice Knepper of Villanova University and were cultured in Dulbecco’s Modified Eagle Medium (DMEM).

### Cell Transfections

MB49, PC3, and MCF-7 cells were separately seeded on twenty-four well plates at a density of 50,000 cells/well in fetal bovine serum-containing media (SCM) twenty-four hours prior to transfection. Polyplexes were prepared by mixing PEI and the luciferase expression plasmid pGL4.50 in a 5:1 w/w ratio with a total of 200 ng DNA/well, then incubating the mixture for twenty minutes at room temperature. Meanwhile, the SCM in each well was aspirated and replaced with serum-free media (SFM). Polyplexes were simultaneously added to each well with inhibitors at a range of concentrations (1 nM - 100 μM final concentrations), while control wells received polyplex without inhibitor. Cells were then incubated for an additional six hours at 37°C in SFM, after which time the media was exchanged again with fresh SCM containing the inhibitors. Cells were then incubated for forty-eight hours before transgene (luciferase) expression was measured with a luminescence assay. Raw luminescence values for each sample were then normalized to the luminescence of the polyplex control (with no drug added) to obtain the relative luminescence displayed in Figure 2.

**Figure 2.**
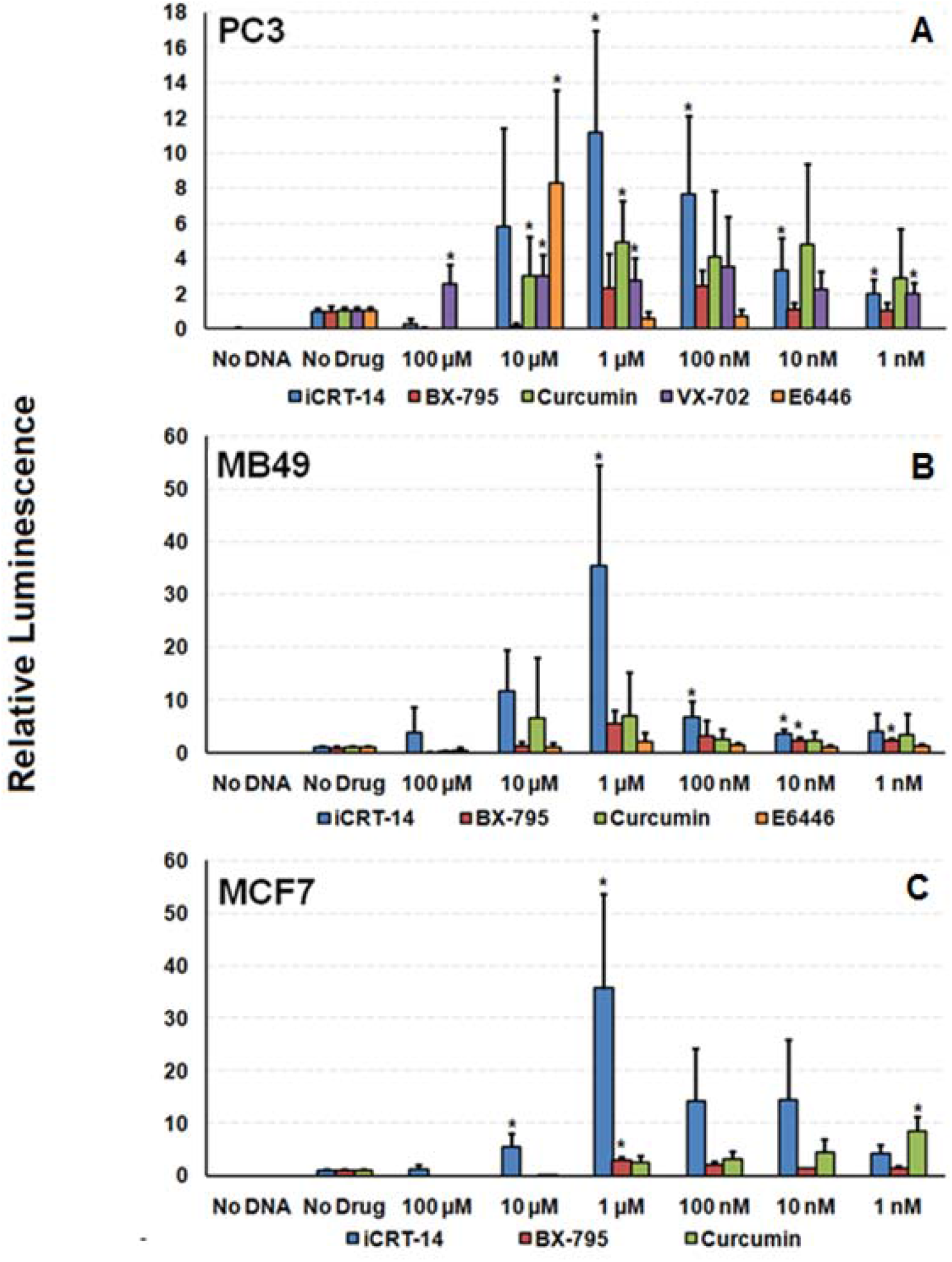
Effects of inhibitors on transgene (luciferase) expression in PC3 **(A)**, MB49 **(B)**, and MCF-7 **(C)** cell lines. Significance was determined with one tail student T tests for values greater than the “No Drug” control. (*p < 0.05) n ≥ 3 for each cell line tested.

### MTT Assay

In addition to transgene expression, the proliferation of the transfected cells was also measured with the MTT (3-(4,5-dimethylthiazol-2-yl)-2,4-dipehnyltetrazolium bromide) assay to quantify the effects of each inhibitor on cell proliferation. Two days after transfection, 10 μL of 5.0 mg/mL MTT was added directly to the cells and incubated for two hours at 37°C. During that time, living cells reduced the yellow MTT dye into purple formazan. 150 μL of DMSO was then added to each well to lyse the cells and dissolve the formazan, which was quantified by measuring the sample’s absorbance at 590 nm. Relative proliferation was then calculated by dividing the absorbance of each sample by the absorbance of the live cell control, in which no polyplex or drug was added.

### ELISA: Cytokine Quantification

To confirm the inhibition of the IIR by SMIs, an enzyme-linked immunosorbent assay (ELISA) was conducted to quantify IL-6 production in PC3 cells at six and twenty-four hours after the addition of polyplexes. In these experiments, cells were transfected as usual, but the supernatant SFM/SCM was collected, centrifuged at 1,000 g for ten minutes and stored at −72°C until the ELISA was performed. The ELISA procedure was conducted according to the manufacturers (R&D Systems) instructions, in which the presence of IL-6 is detected via a colorimetric assay. A standard calibration curve using provided recombinant IL-6 was prepared to estimate IL-6 concentrations.

## Results and Discussion

### Effects of Inhibitors on Transgene (Luciferase) Expression

The effects of each inhibitor on transgene expression in PC3, MCF-7, and MB49 cell lines are shown in Figure 2. In prostate cancer (PC3) cells, 10 μM E6446 and 1 μM iCRT-14 provided the highest significant increases in relative luminescence (8.3 ± 5.3-fold and 11.2 ± 5.8-fold, respectively). Luciferase expression was also modestly enhanced by a factor of 4.9 ± 2.4 by 1 μM Curcumin and 3.5 ± 2.9-fold by VX-702. In contrast, while previous studies have shown that the TBK-1 inhibitor BX-795 increases transgene expression in NK cells, we observed no significant enhancement of luciferase expression by BX-795 in PC3 cells.^20^ Likewise, inhibitors of MNK1 (CGP 57380) and IRAK1/4 (IRAK1/4i) also showed no significant enhancement (data not shown).^29^

Similar trends were also observed in the murine bladder cancer cell line (MB49), with 1 μM iCRT-14 significantly increasing transgene expression, but at a much higher magnitude (35.5 ±19-fold). BX-795 (1 μM) also provided a modest enhancement in MB49 cells (2.5 ± 0.2-fold). In contrast, E6446 showed a relatively lower level of enhancement (2.2 ± 1.5-fold) at a concentration of 1 μM (instead of the 10 μM optimum observed in PC3 cells). VX-702 was also tested in MB49 cells, but no significant enhancement was observed (data not shown).

In the final round of transfections, iCRT-14, BX-795, and curcumin were also tested in a breast cancer cell line (MCF-7). In this second human cell line, 1 μM iCRT-14 continued to significantly increase relative luminescence by 36 ± 18-fold. Curcumin also significantly increased luciferase expression relative to the polyplex control by a factor of 8.5 ± 2.8-fold, but at a much lower optimum concentration (1 nM) than previously observed in the other cell lines (1 μM). Finally, 1μM BX-795 also provided a relatively modest enhancement of 2.9 ± 0.4-fold. This result is similar to the work conducted by Sutlu et al, in which BX-795 provided a 3.8-fold enhancement of transgene expression in NK cells.^26^

Overall, iCRT-14 provided the highest enhancement of transgene expression across all cell lines tested. ICRT-14 belongs to a class of molecules called thiazolidinediones, which have been shown to down regulate the Wnt/β-catenin pathway by directly binding to β-catenin and preventing β-catenin-TCF4 interaction.^25,30^ In addition, iCRT-14 may also disrupt the activation of β-catenin by the DNA sensor LRRFIP1 (see Figure 1).^31,32^ In either case, activation of β-catenin allows it to bind IRF3 and p300 acetyltransferase.^31^ This complex then activates specific host genes (e.g. Type I interferons such as IFN-β) via histone hyperacetylation at specific loci (e.g. *Ifnb1* promoter).^33,34^ However, inhibition of β-catenin by iCRT-14 may increase transgene expression by preventing expression of those host genes.

### Effects of Inhibitors on Cell Proliferation

While the luminescence assays clearly demonstrate the positive effects of each inhibitor on transgene expression, it is also important to consider the potential toxicity of each drug. Figure 3 shows the effects of each inhibitor on cell proliferation (viability) in PC3, MCF-7, and MB49 cells. Overall, the polyplex alone with no additional inhibitors consistently decreased cell viability to 80% in all cell lines, which is similar to previously reported results with PEI.^35^ In PC3 cells, most of the inhibitors exhibited anti-proliferative effects above 1 μM (except VX-702). However, the optimum concentrations of iCRT-14 and Curcumin (1 μM for both) showed no significant effect on cell proliferation. The optimum concentration of E6446 (10 μM) did significantly decrease proliferation to 15% of the control, but lower concentrations had no toxic effect. Each drug showed similar trends in the MB49 and MCF-7 cell lines, with high toxicity at the upper concentrations but no significant effects on cell proliferation at their optimum concentrations for transgene enhancement. However, there were a few notable differences. First of all, the BX-795 significantly decreased cell proliferation to 25% of the live cell control at its optimum concentration (10 μM) in MCF-7 cells. E6446 also significantly decreased cell proliferation at its optimum concentration in MB49 cells (1 μM), but to a much lesser extent (70% viability). In contrast, curcumin significantly increased proliferation relative to the polyplex control at a concentration of 1 μM in MB49 cells (a similar increase was also observed in PC3 cells, but the effect was not significant).

**Figure 3.**
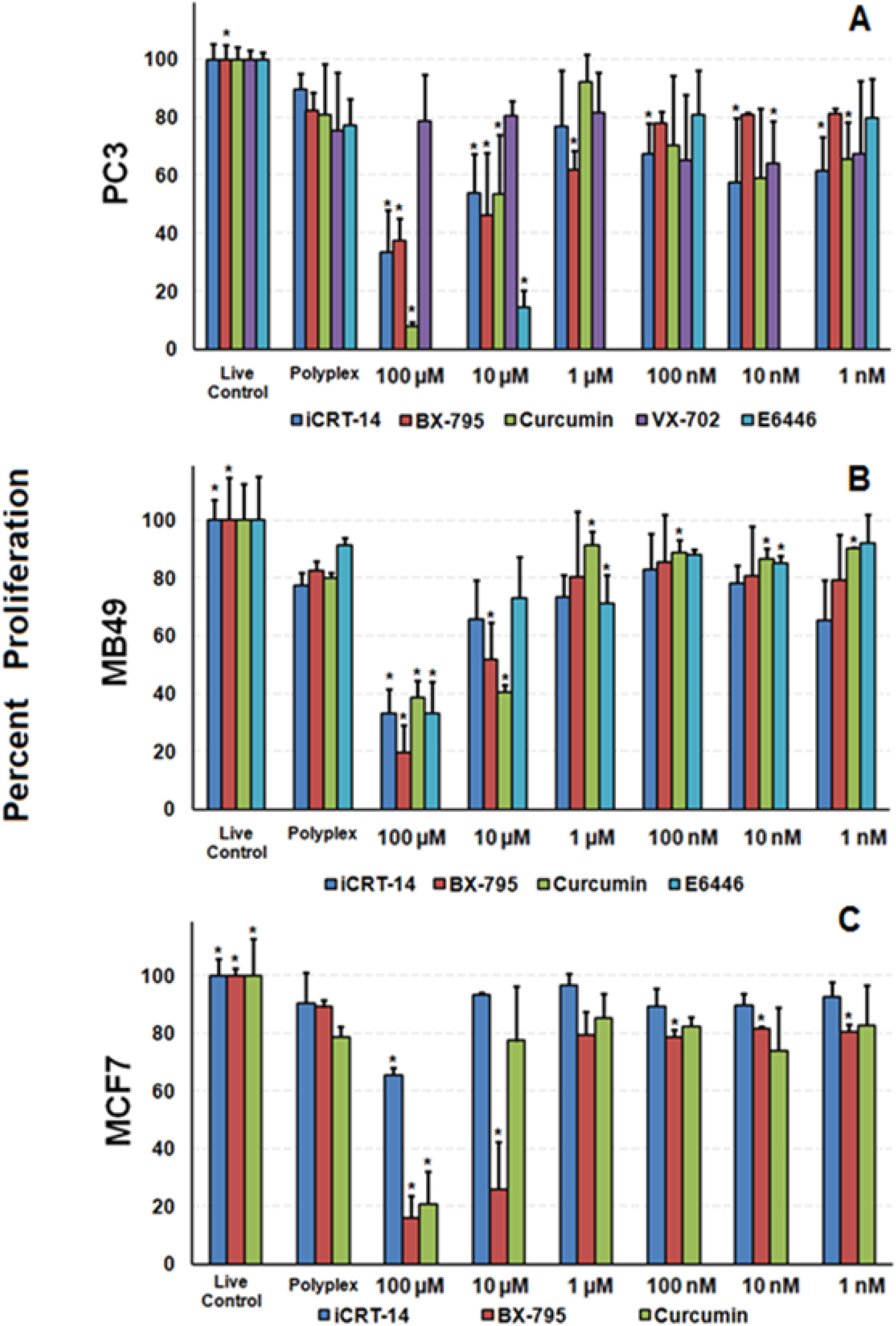
Effects of inhibitors on cellular proliferation in PC3 **(A)**, MB49 **(B)**, and MCF-7 **(C)** cell lines. Significance was determined with one tail student T tests for values greater or less than the polyplex control. (*p < 0.05) n ≥ 3 for each cell line tested.

### Effects of Inhibitors on Inflammatory Cytokine (IL-6) Expression

To confirm that the enhancement observed with each drug was due to inhibition of the innate immune response instead of some other off-target effect, the effect of the inhibitors on inflammatory cytokine (IL-6) expression was also measured in PC3 cells. Specifically, IL-6 was chosen as a marker of IIR activation since it has been previously shown that transfection of plasmid DNA increases IL-6 expression by a factor of 7-12 fold in HeLa cells.^15^ As shown in Figure 4, IL-6 expression was quantified in PC3 cells at six hours post transfection (while the cells were still exposed to serum-free media containing polyplex) and at twenty-four hours post transfection (while the cells were in SCM with inhibitors, but no polyplex present). First of all, it is interesting to note that IL-6 expression in the control samples (only cells; no drug or plasmid) increased an order of magnitude (40 to 400 pg/mL) after the media was changed from SFM to SCM. This motivated us to assay sterile SCM (without cells) for IL-6, but no IL-6 was detected (data not shown). Therefore, this increase may reflect an inflammatory response to components in the fetal bovine serum of SCM. Nonetheless, addition of polyplex to the cells did induce a ~50% increase in IL-6 levels at both time points.

**Figure 4.**
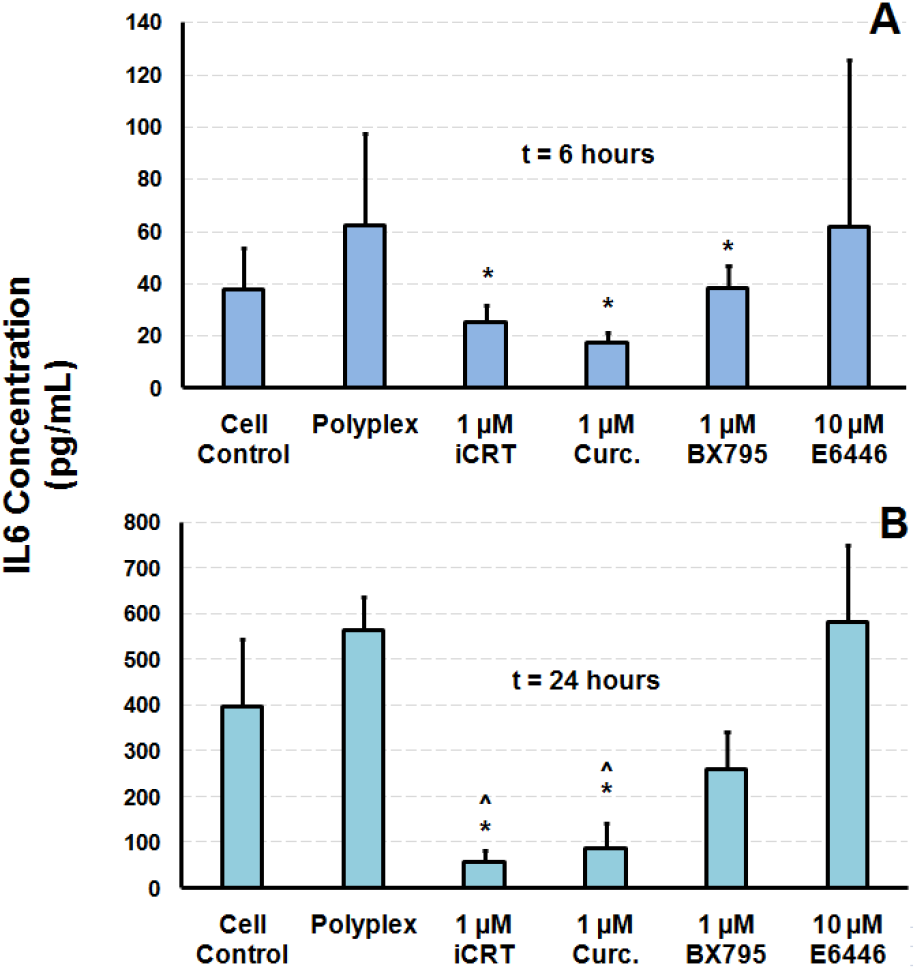
IL-6 expression in the presence of iCRT-14, curcumin, BX-795, and E6446 at six and twenty-four hours post transfection in PC3 cells. Cell controls lacked drug and polyplex. **(A)** Samples at six-hour time points were taken in serum-free media, while samples at **(B)** Twenty-four hour time points were taken in serum-containing media. One tail student T tests were performed with respect to polyplex controls and cell controls (*p < 0.05 for IL-6 concentrations significantly lower than the polyplex control and ^^^p < 0.05 for IL-6 concentrations significantly lower than the cell control).

Compared to the polyplex control, both 1 μM iCRT-14 and 1 μM curcumin significantly decreased IL-6 levels by 60-70% at the six-hour time point relative to the polyplex control. BX-795 (1 μM) also significantly reduced IL-6 expression at the six-hour time point, but this effect was not significant after twenty-four hours. In contrast, the effects of iCRT-14 and curcumin were even more pronounced (80-90% decrease in IL-6 levels) at twenty-four hours post transfection in SCM. It is also interesting to note that both iCRT-14 and curcumin significantly reduced IL-6 production relative to the live cell control to which polyplex was not added. Therefore, iCRT-14 and curcumin may be able to prevent inflammation due to plasmid DNA and the transition to serum-containing media. Similar effects on IL-6 expression have also been observed in other studies involving these inhibitors and pathways. For example, Wnt/β-catenin activation has been shown to increase IL-6 production.^36,37^ Curcumin and BX-795 have also been shown to directly decrease IL-6 production.^38,39^

While E6446 showed significant enhancement of transgene expression, it had no significant effect on IL-6 levels. This result was unexpected, since previous studies have shown that E6446 lowers IL-6 production in dendritic cells that have been exposed to agonists such as unmethylated CpG DNA, RNA oligonucleotides, and lipopolysaccharides (LPS).^23^ However, Obonyo et al showed that TLR9 was not required for the production of IL-6 in bone marrow-derived macrophages upon exposure to the bacterium *H. pylori*. Therefore, inhibition of TLR9 by E6446 alone may not be sufficient to suppress IL-6 production, since the downstream intermediate MyD88 may be activated by DHX9/DHX36 (see Figure 1) to produce IL-6.^40^

The results in Figure 4 and a previous study by Holl et al show that IL-6 is upregulated in response to plasmid DNA.^41^ Once IL-6 is secreted, it may bind to cognate receptors on other cells to activate a JAK/STAT signaling cascade that can activate a many different genes, including transcription factors (JunB, c-Fos, and IRF-1) that further propagate the innate immune response. Interestingly, it has also been shown that inhibition of JAK/STAT signaling with the JAK inhibitor AG-490 significantly increases luciferase expression in PC3 cells.^42^ Altogether, these results show that plasmid DNA activates an innate immune response that increases IL-6 expression and hinders transgene expression, but inhibition of different parts of this pathway can restore transgene expression.

### Synergistic Effects of Inhibitors on Transgene (Luciferase) Expression

Since each of the inhibitors target different components of the highly redundant innate immune response, we hypothesized that combining multiple inhibitors would further increase transgene expression. To test this hypothesis, the three lead inhibitors (10 μME6446, 1 μM iCRT-14, and 1 μM curcumin) were combined in pairs and triplicate in PC3 cells. Figure 5 shows that the combination of E6446 and iCRT-14 seemed to synergistically increase transgene expression relative to the individual inhibitors, but the difference was not statistically significant due to large variations between experiments. Likewise, all other combinations did not significantly enhance expression and the combination of all three inhibitors instead decreased transgene expression significantly, perhaps due to their combined toxicity (see Figure 5b). Interestingly, some of the combinations did significantly alter cell proliferation. For example, while curcumin showed almost no toxicity and E6446 was highly toxic, combining the two drugs appeared to lead to an intermediate toxicity. Similar effects were also observed for the combination of iCRT-14 and E6446.

**Figure 5.**
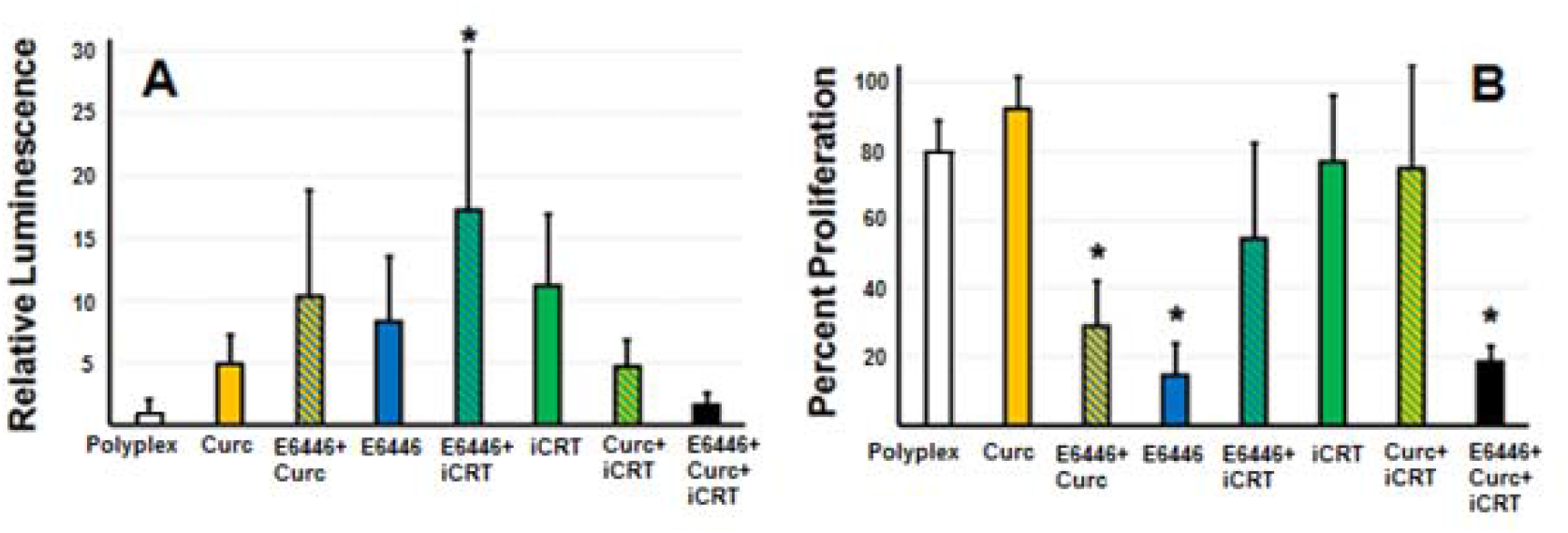
Combinatorial effects of inhibitors on **(A)** transgene (luciferase) expression and **(B)** cellular proliferation. One tail student T tests were performed with respect to individual drugs for luciferase expression and polyplex controls for cellular proliferation (*p < 0.05 for samples with significantly greater luminescence or significantly lower percent proliferation relative to the polyplex control).

## Conclusion

Overall, this study shows that inhibition of the innate immune response can significantly improve transgene expression. Specifically, the β-catenin inhibitor iCRT-14 is able to potently increase transgene expression in a variety of cancer cell lines by decreasing IL-6 expression without a significant decrease in cell proliferation at optimal concentrations. However, since systemic inhibition of the IIR may make a patient vulnerable to infection, future studies may be needed to investigate alternative localized mechanisms of inhibition (e.g. gene silencing/knockout).

## Acknowledgements

The authors would like to gratefully acknowledge Dr. Christine Voekel-Johnson and Dr. Janice Knepper for cell lines and advice. The authors would also like to acknowledge Dr. Noelle Comolli for advice on ELISA assays and Eisai Co. for generously providing the small molecule inhibitor E6446.

